# Bridging the Gap: Exploring Neural Mechanisms of Facial Repetition Suppression through EEG and DCNNs

**DOI:** 10.1101/2023.01.02.522298

**Authors:** Zitong Lu, Yixuan Ku

**Affiliations:** Department of Psychology, The Ohio State University, Columbus, OH, USA; Guangdong Provincial Key Laboratory of Brain Function and Disease, Center for Brain and Mental Well-being, Department of Psychology, Sun Yat-sen University, Guangzhou, China; Peng Cheng Laboratory, Shenzhen, China

**Author notes:** **Authorship contribution statement**Z.L. and Y.K. designed research; Z.L. performed research; Y.K. supervised research; Z.L. and Y.K. wrote the paper. **Declaration of Competing Interest** The authors declare that they have no known competing financial interests or personal relationships that could have appeared to influence the work reported in this paper. **Data availability** All analysis scripts are available online at osf.io/unhzm.

**Keywords:** repetition suppression, reverse engineering, EEG, DCNN, RSA

## Abstract

Facial repetition suppression, a well-studied phenomenon characterized by decreased neural responses to repeated faces in visual cortices, remains a subject of ongoing debate regarding its underlying neural mechanisms. Recent advancements have seen deep convolutional neural networks (DCNNs) achieve human-level performance in face recognition. In our present study, we sought to compare brain activation patterns derived from human electroencephalogram (EEG) data with those generated by DCNNs. Employing reverse engineering techniques, we aimed to provide a novel perspective on the neural mechanisms underlying facial repetition suppression. Our approach involved employing brain decoding methods to investigate how representations of faces change with their familiarity in the human brain. Subsequently, we constructed two models for repetition suppression within DCNNs: the Fatigue model, which posits that stronger activation leads to greater suppression, and the Sharpening model, which suggests that weaker activation results in more pronounced suppression. To elucidate the neural mechanisms at play, we conducted cross-modal representational similarity analysis (RSA) comparisons between human EEG signals and DCNN activations. Our results revealed a striking similarity between human brain representations and those of the Fatigue DCNN, favoring the Fatigue model over the Sharpening hypothesis in explaining the facial repetition suppression effect. These representation analyses, bridging the human brain and DCNNs, offer a promising tool for simulating brain activity and making inferences regarding the neural mechanisms underpinning complex human behaviors.

## Introduction

In our daily experiences, we frequently encounter recurring or similar stimuli alongside new ones. When we repeatedly receive the same or similar information input, our brain’s neural activity tends to diminish compared to the initial exposure, a phenomenon referred to as repetition suppression. Numerous electrophysiological studies have observed that neurons sensitive to visual information in the interior temporal cortex exhibit reduced responses when exposed to repetitive stimuli (Baylis & Rolls, 1987; Kaliukhovich & Vogels, 2011, 2012; Miller et al., 1991; Ringo, 1996; Sawamura et al., 2006; Sobotka & Ringo, 1994). Additionally, research has shown that repeated stimuli can lead to a decrease in the blood oxygenation level-dependent (BOLD) response in functional magnetic resonance imaging (fMRI) studies (Henson & Rugg, 2003). Within the field of face perception, numerous electroencephalogram (EEG) and Magnetoencephalography (MEG) studies have reported various event-related potential (ERP) components associated with facial repetition suppression. These include the N170 (Kloth et al., 2010; Kloth & Schweinberger, 2010; Kovács et al., 2006; Maurer et al., 2008; Mercure et al., 2011; Schweinberger et al., 2007; Walther et al., 2013), P200 (Burkhardt et al., 2010; Kaufmann & Schweinberger, 2012; Latinus & Taylor, 2006; Schulz et al., 2012; Zheng et al., 2012), N250r (Dörr et al., 2011; Herzmann et al., 2004; Pfütze et al., 2002; Schweinberger et al., 1995; Schweinberger & Burton, 2003; Wiese et al., 2013), and N400 (Barrett & Rugg, 1989; Bentin et al., 1985; Rugg, 1985; Schweinberger, 1996; Stevenage et al., 2014). Despite these observations, the precise neuronal mechanism responsible for repetition suppression remains a topic of ongoing debate.

Previous research on facial repetition suppression has predominantly relied on univariate analysis methods, often overlooking the dynamic nature and variances in neural representations. However, in the past decade, cognitive neuroscience has seen a growing trend toward the adoption of multivariate analysis techniques. These include methods like correlation or classification-based Multivariate Pattern Analysis (MVPA) (Cox & Savoy, 2003; Golomb & Kanwisher, 2012; Haxby et al., 2001; Kamitani & Tong, 2005; Norman et al., 2006) and representational similarity analysis (RSA) (Kriegeskorte, Mur, & Bandettini, 2008; Kriegeskorte, Mur, Ruff, et al., 2008). These multivariate analysis tools provide valuable insights into the neural mechanisms underlying complex cognitive processes by capturing the representational patterns through which our brains encode information. RSA, in particular, enables researchers to conduct representational comparisons across diverse modalities. For instance, it allows for the comparison of brain activity patterns with activations in computational models (Cichy et al., 2016; Dobs et al., 2019; Güçlü & van Gerven, 2015; Kuzovkin et al., 2018; Urgen et al., 2019; Xie et al., 2020; Xu & Vaziri-Pashkam, 2021; Yamins et al., 2014). These advanced methods open up new avenues for a deeper understanding of neural processing and encoding in the brain.

Grill-Spector et al., 2006 proposed three potential models (Fatigue, Sharpening, and Facilitation) to explain repetition suppression in neural coding patterns, drawing from a range of studies involving single-cell recordings, fMRI, and EEG/MEG. These models offer different hypotheses about how the brain processes repeated stimuli. The Fatigue model proposes that neurons with stronger initial responses to a stimulus exhibit higher repetition suppression. The Sharpening model suggests that neurons encoding irrelevant features of the stimulus show repetition suppression, leading to a more focused representation. The Facilitation model posits that repetition accelerates stimulus processing, reducing waiting time. To determine which of these models is more likely to underlie facial repetition suppression in human brains, multivariate analysis techniques can be employed.

Recent advances in computer vision have led to the development of deep convolutional neural network (DCNN) models for face recognition (Parkhi et al., 2015; Ranjan et al., 2019; Schroff et al., 2015; Taigman et al., 2014) that have achieved human-level performance (Phillips et al., 2018). In parallel, researchers in cognitive neuroscience and computer science have begun exploring the similarities and differences between human brains and artificial intelligence (AI) models in information processing. Studies that combine brain activity measurements and DCNNs have found that the hierarchical structure of the ventral visual pathway and DCNNs share similar processing representations of visual information (Cichy et al., 2016; Güçlü & van Gerven, 2015; Kietzmann et al., 2019; Yamins et al., 2014). The idea of reverse engineering, where the representation of an AI model can be modified based on theoretical hypotheses to align with the representation of human brains, provides a promising solution to investigate the neural mechanism of repetition suppression. Therefore, we can potentially apply reverse engineering methods to explore the correspondence between the representation of AI models and human neural representations of facial repetition suppression.

Our study aimed to delve into the neural mechanism of facial repetition suppression using innovative computational approaches. Initially, we investigated the dynamic representations of facial information and confirmed the presence of the facial repetition suppression effect in human brains. We accomplished this using a classification-based MVPA method on human EEG data. Subsequently, we employed the concept of reverse engineering to develop two potential repetition suppression models, namely the Fatigue and Sharpening models. These models were used to manipulate the activation of a DCNN. We then conducted cross-modal RSA between human brain activity and the activations in the modified DCNNs. Our findings strongly suggest that the fatigue mechanism is the more plausible neural mechanism underlying facial repetition suppression in the human brain.

## Material and methods

### Data and experimental information

The data utilized in this study were sourced from an EEG dataset focusing on face perception, accessible on OpenNeuro (https://openneuro.org/datasets/ds002718/). This dataset is a comprehensive multi-subject, multi-modal human neuroimaging dataset (Wakeman & Henson, 2015). In our analysis, we specifically utilized EEG data from 18 subjects, which featured 70 valid channels. All subjects participated in a face perception task (Figure 1A) that involved the presentation of 450 grayscale face stimuli. This set comprised 150 familiar faces, 150 unfamiliar faces, and 150 scrambled faces. Each face stimulus was presented twice, with a 50% probability of seeing the same face either immediately or after a delay of 5 to 15 trials. Additional details regarding the experimental design can be found in Wakeman & Henson, 2015.

**Figure 1.**
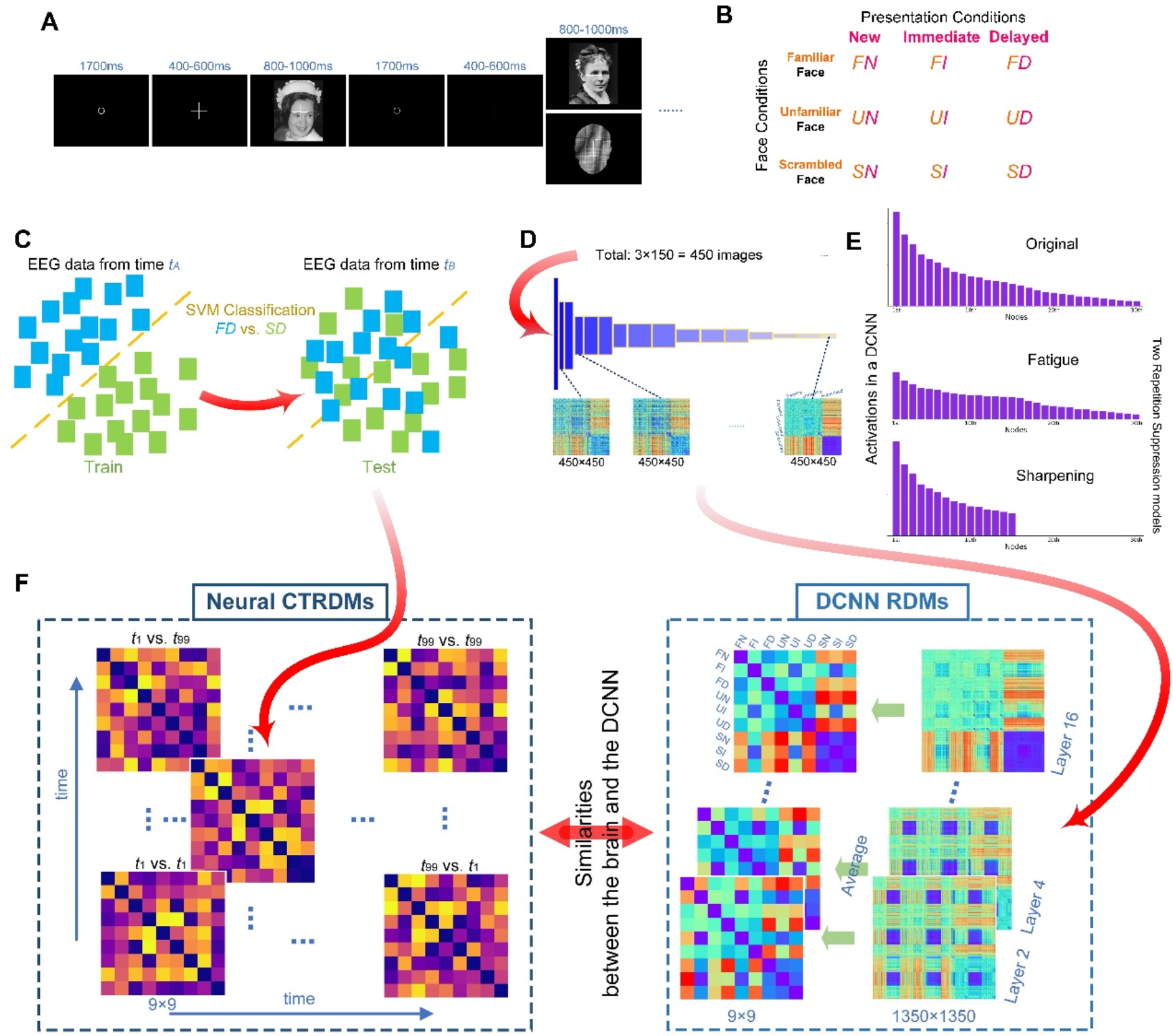
Experimental Procedure and Key Analysis Flow Chart. (A) Overview of the experimental sequences. (B) Illustration of the 9 experimental conditions, consisting of 3 face conditions by 3 presentation conditions. (C) Schematic representation of cross-temporal EEG decoding. (D) Diagram depicting the process of calculating DCNN RDMs (Representational Dissimilarity Matrices). (E) Schematic illustration of the Fatigue and Sharpening repetition suppression models. (F) Flowchart outlining the procedure for cross-modal RSA (Representational Similarity Analysis) comparisons between EEG and DCNNs. Note: The two face images displayed here are from the public domain and are available at https://commons.wikimedia.org for illustrative purposes only. The actual images used during the experiment were described in Wakeman & Henson, 2015.

In our analysis, we labeled the first time a subject saw a specific face as ’new,’ the second time they saw the same face immediately as ’immediate (repetition),’ and the second time they saw the face after some trials as ’delayed (repetition).’ This resulted in a total of 9 conditions (FN, FI, FD, UN, UI, UD, SN, SI, SD) derived from the combination of 3 face conditions (F: familiar, U: unfamiliar, S: scrambled) and 3 presentation conditions (N: new, I: immediate, D: delayed) (Figure 1B).

The EEG data employed in this study had already undergone partial preprocessing and had been resampled to 250 Hz as part of the dataset. We applied band-pass filtering to retain data within the 0.1 to 30 Hz range. To identify and eliminate blinks and eye movements, we utilized independent components analysis (ICA) (Drisdelle et al., 2017; Jung et al., 2000) using EEGLAB (Delorme & Makeig, 2004). Epochs of -500 to 1500ms from stimulus onset were created. No baseline correction was applied.

### Classification-based EEG decoding

#### Time-by-time EEG decoding

Each EEG trial corresponded to two distinct labels: one indicating the face condition (F, U, or S) and the other indicating the presentation condition (N, I, or D). To investigate neural representations across different presentation conditions, we performed nine separate classification-based decoding analyses, aiming to distinguish between them. These analyses encompassed the following condition pairs: FN vs. FI, UN vs. UI, SN vs. SI, FN vs. FD, UN vs. UD, SN vs. SD, FI vs. FD, UI vs. UD, SI vs. SD.

For each of these classification-based decoding analyses, we employed linear Support Vector Machines (SVM). Our approach involved binarizing the trial labels for the two conditions being compared. To reduce the dimensionality of the EEG data, we downsampled it by averaging every five time-points. The original 500 time-points spanning from -500 to 1500ms were compressed into 100 time-points. This yielded a label vector for each subject, containing labels for all trials, as well as a three-dimensional matrix with dimensions for time, trial, and channels, facilitating time-by-time classification.

In our process, we randomized the order of trials and then averaged the data every five trials. The classifier was trained and tested separately for each time-point. Specifically, we randomly selected 2/3 of the trials for training and used the remaining 1/3 for testing in each iteration. This entire sequence of random shuffling, averaging, classification training, and testing was repeated 100 times for each time-point. We repeated these steps for each subject and for all nine classification condition pairs. To obtain more reliable time-by-time decoding accuracies, we averaged the classification accuracies across all iterations. This entire process was replicated for the 18 subjects in our study.

#### Cross-temporal EEG decoding

Additionally, we carried out cross-temporal EEG decoding, which represents an extension of the time-by-time decoding approach. The fundamental idea behind cross-temporal decoding is to train the classifier on data at one specific time-point and then test it on data from other time-points to assess whether the encoding patterns of the information of interest remain consistent across different times.

Building upon the time-by-time decoding methodology described earlier, we proceeded to test each classifier on data from all 100 time-points, including the time-point that was used for training (see Figure 1C for a graphical representation). Similar to the previous approach, we obtained the final decoding accuracies by averaging results across 100 iterations. Consequently, each subject generated nine temporal generalization matrices for cross-temporal decoding, each corresponding to one of the nine classification condition pairs. All the EEG decoding processes described above were executed using the NeuroRA toolbox (Lu & Ku, 2020).

### DCNN models

In this study, we employed a DCNN model commonly utilized in the field of face recognition known as VGG-Face (Parkhi et al., 2015) (Supplementary Figure 1A). VGG-Face was pretrained on a dataset consisting of 2622 unique identities, each with 1000 face images per person. The model exhibited impressive test accuracies, achieving 97.27% on the IFW dataset and 92.8% on the YouTube Faces dataset. VGG-Face essentially followed the structure of a VGG-16 model, comprising 13 convolutional layers and 3 fully connected layers. In our study, we utilized the VGG-Face model as a DCNN for extracting facial features. For the sake of comparison, we also incorporated an additional VGG-16 model that was left untrained and initialized with random weights. This untrained VGG model served as a DCNN model that did not possess any learned facial features.

### RSA

#### EEG RDMs

Given that we had 3 face conditions and 3 presentation conditions, we utilized EEG classification-based decoding accuracy as the dissimilarity metric to build a set of 9x9 neural Representational Dissimilarity Matrices (RDMs). Additionally, we generated Cross-Temporal RDMs (CTRDMs) instead of traditional RDMs to conduct Cross-Temporal Representational Similarity Analysis (CTRSA) (Lu, 2020). In CTRSA, we incorporated cross-temporal decoding accuracy instead of traditional RSA. Here’s how it worked:

As illustrated in Figure 1C, we trained a Support Vector Machine (SVM) classifier on EEG data for the FD vs. SD conditions at timepoint *tA* and then tested it on data for the FD vs. SD conditions at timepoint *tB*. The resulting test accuracy was employed as the dissimilarity index between the FD condition at *tA* and the SD condition at *tB* in a *tA* -> *tB* CTRDM for a particular subject. Due to the directionality from the training time to the testing time, the CTRDM for *tA* -> *tB* was distinct from the CTRDM for *tB* -> *tA*. The former was constructed using data from *tA* for training and *tB* for testing, while the latter was constructed using data from *tB* for training and *tA* for testing. As we couldn’t perform classification between two identical conditions, the diagonal values in the decoding-based CTRDMs were consistently set to 0. By following this approach, we established a CTRDM for each pair of directed timepoints (from one timepoint to another) as described above. Consequently, we generated a total of 100 (timepoints) x 100 (timepoints) CTRDMs based on cross-temporal EEG decoding. The EEG RDMs section was executed using the NeuroRA toolbox (Lu & Ku, 2020).

#### DCNN RDMs

To handle the substantial number of nodes in each layer of the VGG-16 model, we initially applied Principal Component Analysis (PCA) to reduce the feature dimension. For instance, the second layer encompassed 64 112x112 feature maps, equating to a total of 802,816 nodes. For each image fed into the VGG-16 model, the layer 2 activation could be represented as a 1x802,816 vector. We conducted PCA on these vectors and organized the principal components in descending order of their contribution rates. We retained principal components that collectively contributed to over 95% of the variance, discarding the rest. This process effectively reduced the feature dimension of each layer. As depicted in Supplementary Figure 1B, the dimensionality of layer 2 was trimmed to 307 after PCA. In a similar manner, we performed dimension reduction for all layers in both the VGG-Face and untrained VGG models. This dimension reduction step was implemented using the Scikit-learn toolkit (Pedregosa et al., 2011).

To process the images, we fed each one into both the VGG-Face and untrained VGG models, extracting activation vectors for every even layer (e.g., layer 2, 4, …, and layer 16) after dimension reduction. This resulted in 450 activation vectors for each layer, corresponding to the 450 images in our dataset. To manage the computational load, we calculated the Pearson correlation coefficient (r) between the activation vectors for any two images within each layer. We then used 1 minus the correlation coefficient (1 - r) as the dissimilarity index. For a given layer, we constructed a 450x450 Representational Dissimilarity Matrix (RDM) in the order of 150 familiar faces, 150 unfamiliar faces, and 150 scrambled faces. This procedure allowed us to obtain DCNN RDMs for all even layers in both the VGG-Face and untrained VGG models. The calculation of DCNN RDMs was carried out using the NeuroRA toolbox. (Lu & Ku, 2020).

#### Repetition suppression simulations in DCNN

To investigate the neural mechanism of facial repetition suppression using “reverse engineering,” we developed two possible neuronal-level models: the Fatigue model and the Sharpening model. These models were designed to simulate how the neural responses in a deep convolutional neural network (DCNN) change under facial repetition suppression conditions. We set the activation vector of a face image *p* at layer *i* of a DCNN (before PCA) as *A* = (*a*_1_, *a*_2_, … …, *a*_n_), where theactivation values were ordered in descending order (*a*_1_, *a*_2_> *a*_3_, …, *a_m_*_-1_> *a*_m_ = *a_m_*+1 = ⋯ = *a_n_* = 0), which meant that there were *n-m* nodes with nonzero activation value and m nodes with activation value of zero).

For Fatigue model, we assumed that the activation of the node with higher response to face stimulus was weaken under repetition suppression condition. The activation of the node with low response remained unchanged, but the node with higher activation had more attenuations. Thus, if the face image *p* was viewed repeatedly, the new activation vector obtained based on Fatigue model would be *A_F_*= (*a_F_*_1_, *a_F_*_2_, … …, *a_F_*_n_), and its internal activations were:

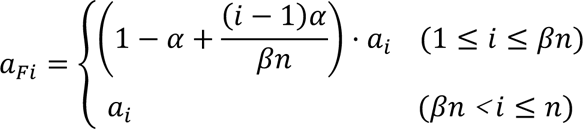

where *α* was the maximum fatigue coefficient, and *β* was the proportion of the nodes that would be attenuated. Thus, the first *βn* nodes would be attenuated when the same face image was viewed repeatedly, and the nodes from the strongest one to the *βn*th one would be weakened in proportion from *α* to *α*/(*βn*).

For Sharpening model, we assumed that the nodes with lower response to face stimulus were no longer activated under repetition suppression condition, and the nodes with higher response kept same activations. Thus, the new activation vector obtained based on Sharpening model would be *A*_*_ = (*a_s1_*, *a_s2_*, … …, *a_sn_*), and its internal activations were:

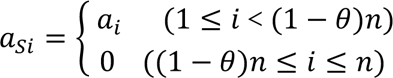

where *θ* was the proportion of the nodes which would be not activated. Thus, the last *θn* nodes’ activations would become zero when the same face image was viewed repeatedly.

Figure 1E is a schematic diagram of how these two repetition suppression models simulated in DCNNs. Here, Figure 1E shows the original activations of 30 active nodes and the activations under repetition suppression condition based on Fatigue model (*α* =0.5, *β* =0.5) and Sharpening model (*θ* =0.5) respectively. For all 450 images, we input them into both VGG-Face and untrained VGG respectively, and then we calculated activation vectors of different layers based on different simulation models of repetition suppression.

#### Modified DCNN RDMs

Based on the above two facial repetition suppression models, we set model parameters corresponding to three types of face stimuli here: For Fatigue model, *α* was set to 0.9, which meant that the first 90% nodes with nonzero activation value were set as the nodes with high activation. And we set the maximum fatigue coefficient *β* to 0.5 for immediate repetition condition and 0.05 for delayed repetition condition. For Sharpening model, *θ* was set to 0.5 for immediate repetition condition and 0.05 for delayed repetition condition. Therefore, three activation vectors corresponding to new, immediate, and delayed conditions were calculated from the activation in each layer in DCNNs of each image based on each repetition suppression model. Then, we input these vectors into PCA to get feature vectors after dimension reduction. For each even layer, face image and repetition suppression model, we calculated the dissimilarity (1-Pearson correlation coefficient) between each pair of two feature vectors and got two 1350×1350 RDMs for VGG-Face and untrained VGG.

#### RSA between EEG and DCNNs

For VGG-Face and untrained VGG, 8 1350×1350 RDMs corresponding to 8 even layers were obtained. For EEG, each subject corresponded 100 time-by-time 9×9 RDMs and 100×100 cross-temporal 9×9 CTRDMs. To establish the connection between DCNNs and human brains, we averaged the cells under 9 conditions respectively (FN, FI, FD, UN, UI, UD, SN, SI, SD) in 1350×1350 RDMs to get compressed 9×9 DCNN RDMs. Then, we calculated the partial Spearman correlation coefficient as the similarity between neural (CT)RDMs and DCNN RDMs for each time- point. To obtain the similarity between the representation of face information that a DCNN learned for face recognition and human brains, we calculated the valid representational similarity *S*_*valid*_ below:

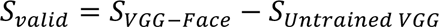

where *S_vgg-Face_* was the representational similarity between VGG-Face and neural activity, and *S_UntraibedVGG_* was the representational similarity between untrained VGG and neural activity. Here, we applied untrained VGG’s representations as a baseline that there was not enough face-specific information and calculated valid similarities for two repetition suppression models, respectively. To get face-specific repetition suppression mechanism, the pre-train VGG-Face trained on face recognition task would have stronger similarity with human brains than untrained VGG.

### Statistical analysis

For the classification-based decoding results, we assessed whether neural representations in the brain encoded information at specific time-points. We assumed that if this was the case, it would be possible to linearly classify between two conditions, resulting in decoding accuracy greater than chance, which is 50%.

Regarding the RSA results, we aimed to determine whether the valid similarity between conditions was significantly greater than zero. If the valid similarity values were either zero or less than zero, it would suggest that the corresponding repetition suppression mechanism was not specific to face information.

To compare the decoding accuracy to chance and assess whether valid similarity differed significantly from zero at each time-point, while also controlling for multiple comparisons, we employed cluster-based permutation tests. Here’s a step-by-step outline of the procedure: (1) Calculate t-values for each time-point and identify significant clusters; (2) Compute the clustering statistic as the sum of t-values within each cluster; (3) Perform 5000 permutations to establish the maximum permutation cluster statistic; (4)Assign p-values to each cluster in the actual decoding accuracies or similarities dataset by comparing their cluster statistic to the permutation distribution. This approach helps us determine the statistical significance of the observed decoding accuracy and similarity values while accounting for multiple comparisons.

### Code availability

All analysis scripts are available online at osf.io/unhzm.

## Results

### Facial repetition suppression in human brains

The classification-based EEG decoding results for the three presentation conditions are presented in Figure 2. Time-by-time decoding results are illustrated in Figure 2A. (1) New vs. Immediate Decoding: Decoding accuracies for both familiar and unfamiliar faces were consistently above chance levels from 200ms to 1500ms. Similarly, decoding accuracies for scrambled faces exceeded chance levels from 360ms to 1500ms. Notably, during specific time intervals, decoding accuracies for familiar faces surpassed those for unfamiliar faces: from 620ms to 800ms, 880ms to 1100ms, and 1120ms to 1220ms. Additionally, decoding accuracies for familiar faces were consistently superior to those for scrambled faces from 240ms to 1500ms. Furthermore, decoding accuracies for unfamiliar faces were significantly better than those for scrambled faces from 240ms to 960ms.

**Figure 2.**
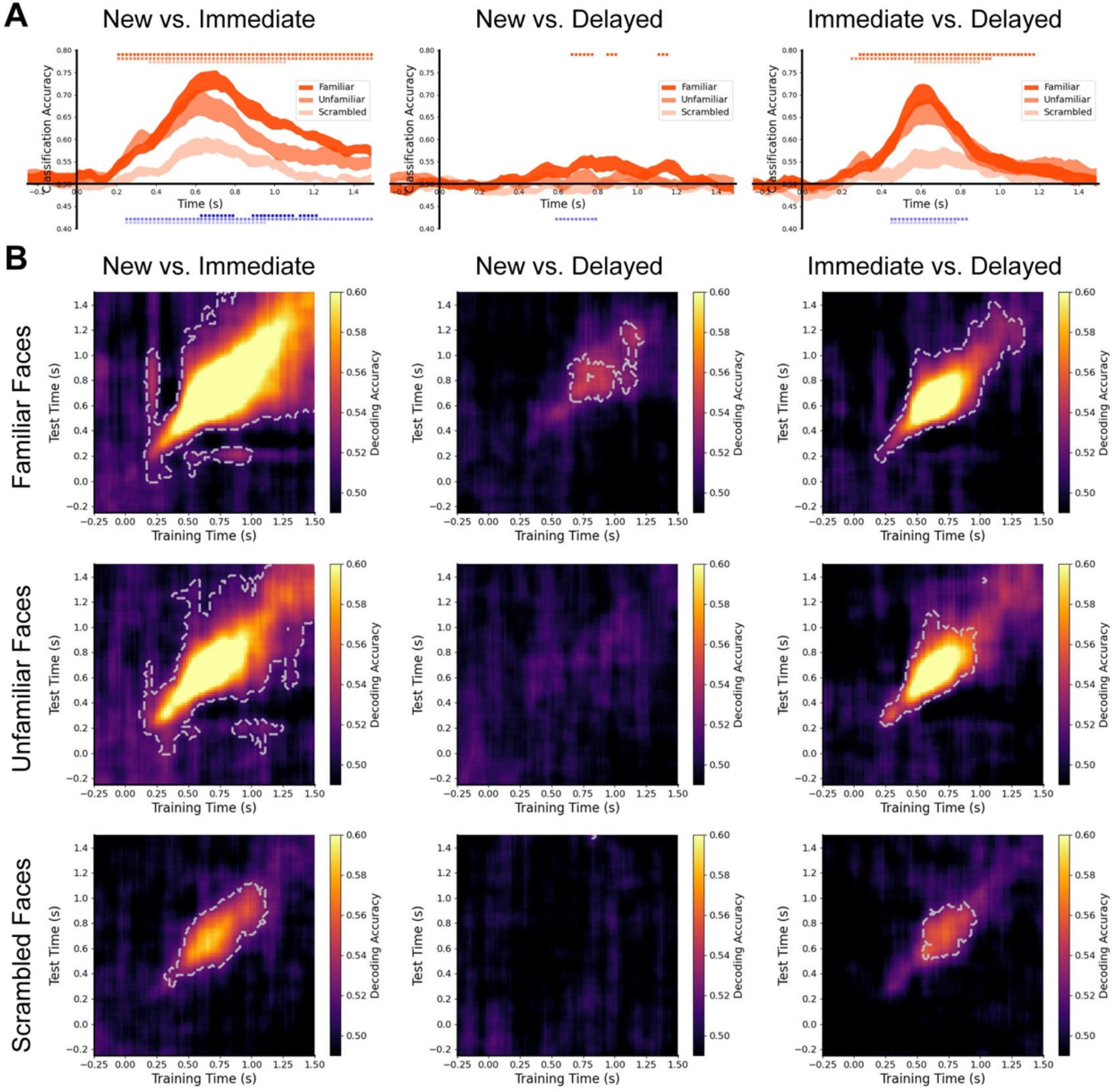
Temporal facial repetition suppression in human brains. (A) Time-by- time decoding results. The top of each plot is adorned with color-coded small squares indicating p<0.01 (cluster-based permutation test) of decoding accuracy significantly greater than chance (ranging from dark to light orange for familiar, unfamiliar, and scrambled faces). The bottom of each plot features color-coded small squares indicating p<0.01 (cluster-based permutation test) of significant differences in decoding accuracy between two face conditions (ranging from dark to light blue for familiar vs. unfamiliar faces, familiar vs. scrambled faces, and unfamiliar vs. scrambled faces). Line width reflects ±SEM. (B) Cross-temporal decoding results. The baseline for classification-based decoding accuracy is 50%. Regions where average accuracy significantly exceeds chance are highlighted with a light grey outline (cluster-based permutation test, p<0.01)..

**Figure 3.**
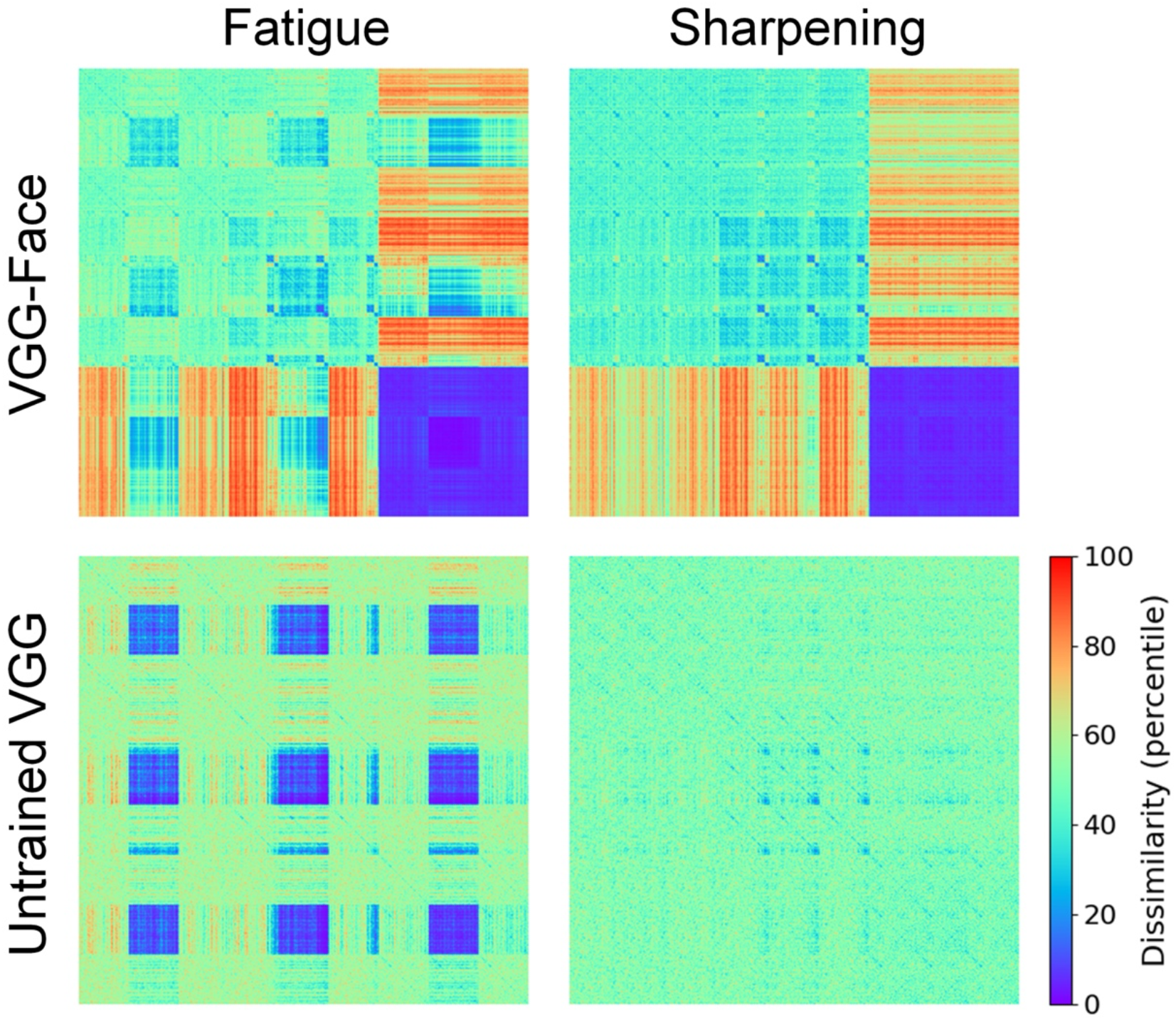
Modified DCNN RDMs of layer 16 based on two different repetition suppression mechanisms.

(2) New vs. Delayed Decoding: Significant decoding accuracies for familiar faces were detected within limited time intervals, specifically from 660ms to 780ms, 860ms to 920ms, and 1100ms to 1160ms. Moreover, during the period from 580ms to 800ms, decoding accuracies for familiar faces were significantly higher than those for scrambled faces.

(3) Immediate vs. Delayed Decoding: Decoding accuracies for familiar faces consistently outperformed chance levels from 280ms to 1180ms. Similarly, decoding accuracies for unfamiliar faces exceeded chance levels from 240ms to 980ms, and those for scrambled faces from 560ms to 900ms. Moreover, during the interval from 440ms to 840ms, decoding accuracies for familiar faces were significantly better than those for scrambled faces, while decoding accuracies for unfamiliar faces surpassed those for scrambled faces from 440ms to 780ms.

Cross-temporal decoding results are displayed in Figure 2B. The most pronounced differences in neural patterns were observed between the New and Immediate conditions. Conversely, there were minimal significant differences between the New and Delayed conditions. Additionally, the differences in neural patterns for familiar faces were more substantial compared to those for unfamiliar faces and scrambled faces.

These results offer valuable insights into the temporal dynamics of neural representations during different presentation conditions, emphasizing the robustness of decoding accuracies in distinguishing between these conditions, particularly in the New vs. Immediate decoding scenario. They shed light on the neural mechanism of facial repetition suppression in human brains. Despite the identical input face images, notable differences were observed in neural representations between new and immediate repetition conditions. This suppression effect diminished as the interval between repetitions increased. Consequently, when the same face image was repeatedly shown after several trials, the neural representation became more similar to that of the initial presentation. Furthermore, familiar faces elicited a stronger repetition suppression effect compared to unfamiliar and scrambled faces.

### Modified representations based on different repetition suppression models in DCNNs

Regarding the DCNNs, we initially extracted features corresponding to the 450 face images from all even layers and computed 450×450 RDMs (Supplementary Figure 1B and 2). To simulate the facial repetition suppression effect in DCNNs, we adjusted neural representations in both VGG-Face and untrained VGG using the Fatigue and Sharpening models, respectively. Figure 2 displays DCNN RDMs of layer 16 derived from DCNN’s internal activations that were modified by the two repetition suppression models. Under the Fatigue model, only the activation of the node with a higher response to the face stimulus exhibited repetition suppression, and nodes with higher activation experienced more suppression. Under the Sharpening model, only the activation of the node with a lower response to the face stimulus exhibited repetition suppression, and nodes with low responses were not activated under repetition conditions. Modified 1350×1350 RDMs for all even layers are presented in Supplementary Figure 3.

### Comparisons between brains and DCNNs revealing a fatigue mechanism

After compressing the DCNN RDMs, we computed the similarity between EEG RDMs and the modified DCNN models, then calculated the difference in cross-modal similarity between VGG-Face and the untrained VGG as the valid similarity. Figure 4A illustrates the valid representational similarity between activations modified by the Fatigue model for all even layers in DCNNs and EEG signals. DCNN representations modified by the Fatigue model exhibited significant representational similarity with human brains. We observed significant valid similarities in many layers (layer 2: 120ms to 640ms, 700ms to 1420ms; layer 4: 120ms to 460ms, 1040ms to 1220ms; layer 10: 480ms to 560ms, 1380ms to 1500ms; layer 12: 260ms to 380ms, 420ms to 560ms, 1380ms to 1500ms; layer 14: 160ms to 1000ms; layer 16: 200ms to 1320ms). However, DCNN representations modified by the Sharpening model showed almost no significant similarity with human brains. We only found significant valid similarities from 640ms to 680ms in layer 8.

**Figure 4.**
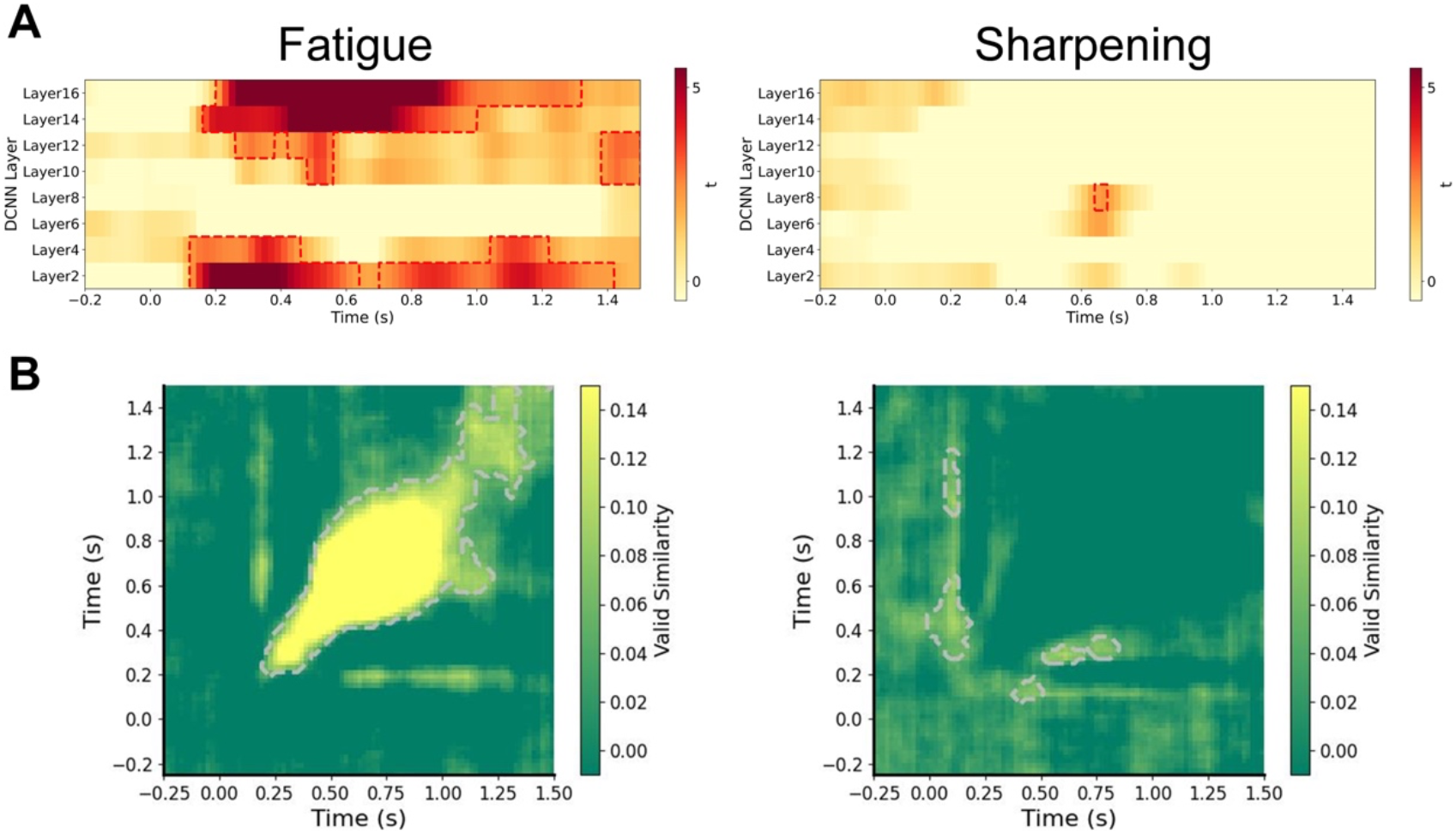
Representational comparison between brain activity and modified DCNN activations. (A) Layer-by-layer temporal valid similarity between EEG and modified DCNNs. (B) Cross-temporal valid similarity between EEG and modified DCNNs on layer 16. The baseline of valid similarity is zero. Outlines indicate significant clusters (cluster-based permutation test, p<0.05).

In detail, Figure 4B presents the cross-temporal valid similarity of layer 16, which is the last layer in the VGG structure and contains the most relevant information for face recognition. When modified by the Fatigue model, the DCNN’s representations exhibited strong and extensive valid similarity with human brains. However, representations of the DCNN modified by the Sharpening model only displayed weak similarities with brain activity.

Cross-modal comparisons suggest that simulating the activation in DCNNs based on a fatigue mechanism, rather than a sharpening mechanism, could induce more similar representations with human brain activities. Therefore, the facial repetition suppression effect in face perception is more likely caused by the fatigue mechanism.

## Discussion

In this study, we employed a unique approach that combines human EEG and deep convolutional neural networks (DCNNs) to delve into the neural mechanism of facial repetition suppression in face perception. We accomplished this through classification-based EEG decoding and cross-modal representational similarity analysis (RSA).

Initially, our EEG-based decoding results provided insights into the temporal dynamics of neural representations across different face presentation conditions. We observed a facial repetition suppression effect that was more pronounced for familiar faces and less so for scrambled faces. This effect decreased as the interval between repeated viewings increased. Notably, our findings revealed distinct neural representations between new and immediately repeated conditions, demonstrating the influence of repetition suppression. Furthermore, the repetition suppression effect was more robust for familiar faces than for unfamiliar or scrambled ones.

Subsequently, we embarked on a reverse engineering endeavor to delve into the mechanisms underlying facial repetition suppression in the human brain. Our approach involved the modification of neural activations in artificial intelligence (AI) models capable of achieving human-level performance in face recognition. Specifically, we honed in on the repetition suppression mechanisms pertaining to face-specific information using the VGG-Face model, a neural network trained on an extensive dataset of facial images for the purpose of face identification. To distill the information relevant to facial features within the deep convolutional neural network (DCNN), we took a unique approach. We contrasted the results obtained from the VGG-Face model with those of an untrained VGG model. This juxtaposition allowed us to extract the valid representational similarity between the trained DCNN and the human brain, focusing on their shared representations of facial stimuli. The outcome of this analysis revealed a significant convergence between the representation of the DCNN model and the neural patterns observed in the human brain when employing the Fatigue model for modification. This alignment was not only observed in the later layers of the DCNN but also extended to the early layers, mirroring the hierarchical structure of the human brain’s visual processing pathway.

Moreover, our cross-temporal analysis illuminated that the similarities between the DCNN model’s representations and the neural patterns observed in the human brain encompassed a broader temporal range. Intriguingly, these similarities weren’t confined solely to the later layers of the DCNN; they extended to the early layers as well. This finding aligns with previous research that has drawn parallels between the hierarchical structure of DCNNs and the visual processing pathway in the human brain (Cichy et al., 2016; Güçlü & van Gerven, 2015). Collectively, these results provide compelling evidence that the attenuation of neural activations in the process of repetition suppression, driven by the fatigue mechanism, occurs not only in neurons processing high-level face features but also in those handling low-level facial information. This underscores the robustness and comprehensiveness of the fatigue-based repetition suppression phenomenon across different neural processing stages, mirroring the multifaceted nature of facial perception in the human brain.

It’s worth noting that while numerous human fMRI studies (Andics et al., 2013; Ewbank et al., 2016; Grotheer et al., 2014; Grotheer & Kovács, 2014;Kovács et al., 2012, 2013; Larsson & Smith, 2012; Mayrhauser et al., 2014) have given support to the idea that repetition suppression is indicative of a reduction in prediction error within the predictive coding framework (Friston, 2005; Summerfield et al., 2008), recent electrophysiological investigations have put forth an alternative perspective. These electrophysiological studies have provided evidence in favor of the fatigue mechanism, positing that it involves bottom-up or local adaptation, as opposed to sharpening or sparseness representations and the predictive coding hypothesis.

However, the collection of human electrophysiological data presents substantial challenges, leading to a scarcity of human neuroimaging studies that could corroborate these findings (Stam et al., 2021). Our present study is innovative in that it introduces state-of-the-art computational methods, combining noninvasive human EEG with DCNNs, to delve into the neural underpinnings of facial repetition suppression. In doing so, we have provided compelling and robust evidence to support the fatigue mechanism as the driving force behind repetition suppression in the context of facial perception.

Furthermore, our study brings to light several areas that warrant further investigation and potential avenues for improvement. Firstly, it’s important to consider whether there are alternative mechanisms underlying facial repetition suppression. While our study examined two primary models, the Fatigue and Sharpening models, it’s conceivable that other mechanisms may be at play. One limitation of pure DCNN models is their lack of inherent temporal processing capability. Future research could explore the inclusion of timing process components, such as recurrent structures (Kietzmann et al., 2019; Spoerer et al., 2017), within DCNN models. This could lead to the discovery of additional mechanisms, including the possibility of a Facilitation mechanism (Grill-Spector et al., 2006) contributing to repetition suppression. Additionally, investigating repetition suppression mechanisms under different conditions and for various types of facial information or tasks could provide valuable insights.

Secondly, the question of whether DCNNs represent the best brain-like models is a valid one. While DCNNs are widely employed in neuroscience, the field of computer vision has witnessed rapid advancements, resulting in the development of new ANN models that demonstrate exceptional task performance and feature space effectiveness. These models include Generative Adversarial Networks (GANs) (Goodfellow et al., 2014), Vision Transformers (ViTs) (Dosovitskiy et al., 2020), unsupervised models(He et al., 2019), and Contrastive Language-Image Pretraining (CLIP) (Radford et al., 2021), and some other brain-inspired models (Li et al., 2021; Zeng & Si, 2021). Additionally, Spiking Neural Networks (SNNs) (Ghosh-Dastidar & Adeli, 2011) and various EEG-based models (Liang et al., 2022; Zhang et al., 2021) have emerged as alternatives. Evaluating which model aligns most closely with the human brain is a complex challenge, and it remains a cutting-edge area of research that garners attention from both computer scientists and neuroscientists. Nevertheless, our study provides a valuable framework for investigating neural mechanisms within the human brain through the use of reverse engineering and cross-modal RSA.

Moreover, our study has certain limitations stemming from the original experimental design of the open dataset we utilized. There exist numerous nuanced facets of facial information that have not been thoroughly explored and deeply analyzed due to these limitations. These unexplored dimensions include variations in hairstyle, skin color, viewpoint (e.g., upright or inverted faces), gender, race, and facial expressions. To gain a more comprehensive understanding of the intricate process of face perception and face repetition suppression, it is imperative that future research incorporates a broader range of experimental stimuli to capture the richness and complexity of facial information processing. From a technical standpoint, our study did not involve the selection of specific EEG channels, and opportunities for channel selection and feature optimization were not fully explored. Implementing engineering techniques, such as channel selection and feature optimization as proposed by Jin et al. (Jin et al., 2019), could yield improvements in decoding accuracy and the quality of representational results. Furthermore, our research primarily relied on EEG data to investigate dynamic neural representations over time. In the future, expanding our investigations to encompass the spatial aspects of neural information processing through fMRI and MEG data could provide valuable insights into how neural representations evolve across different brain regions. This multidimensional approach would offer a more comprehensive perspective on the neural mechanisms underlying facial repetition suppression. In conclusion, this study represents a innovative foray into the realm of neuroscience using state-of-the-art methodologies that fuse artificial intelligence (AI) and neuroimaging techniques. Through the strategic modification of AI model representations, we have endeavored to identify the condition that most closely approximates the neural representations within the human brain. Our exploration of cross-modal representations not only facilitates the unraveling of intricate neuroscience questions that are challenging to address through conventional noninvasive methods such as fMRI or EEG experiments but also holds the potential to inspire advancements in AI models from a neuroscientific perspective. The insights gleaned from this research are poised to be of significant importance for the future development of brain-inspired artificial intelligence. By bridging the gap between AI and neuroscience, this work contributes to a deeper understanding of neural mechanisms and offers invaluable contributions to the ongoing quest for brain-like intelligence.

## Supplementary

**Supplementary Figure 1.**
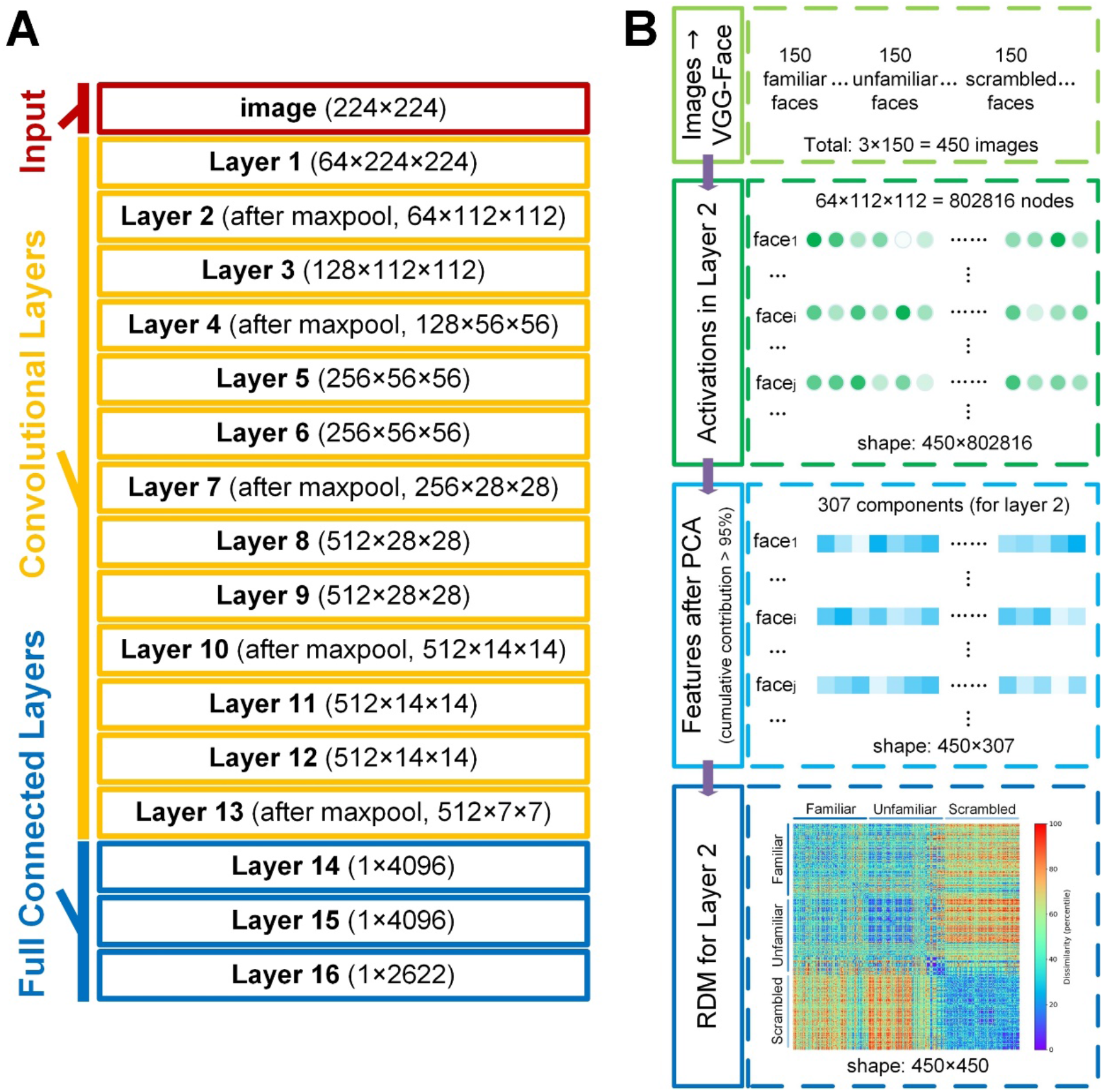
(A) VGG-Face structure and (B) calculation steps of DCNN RDMs.

**Supplementary Figure 2.**
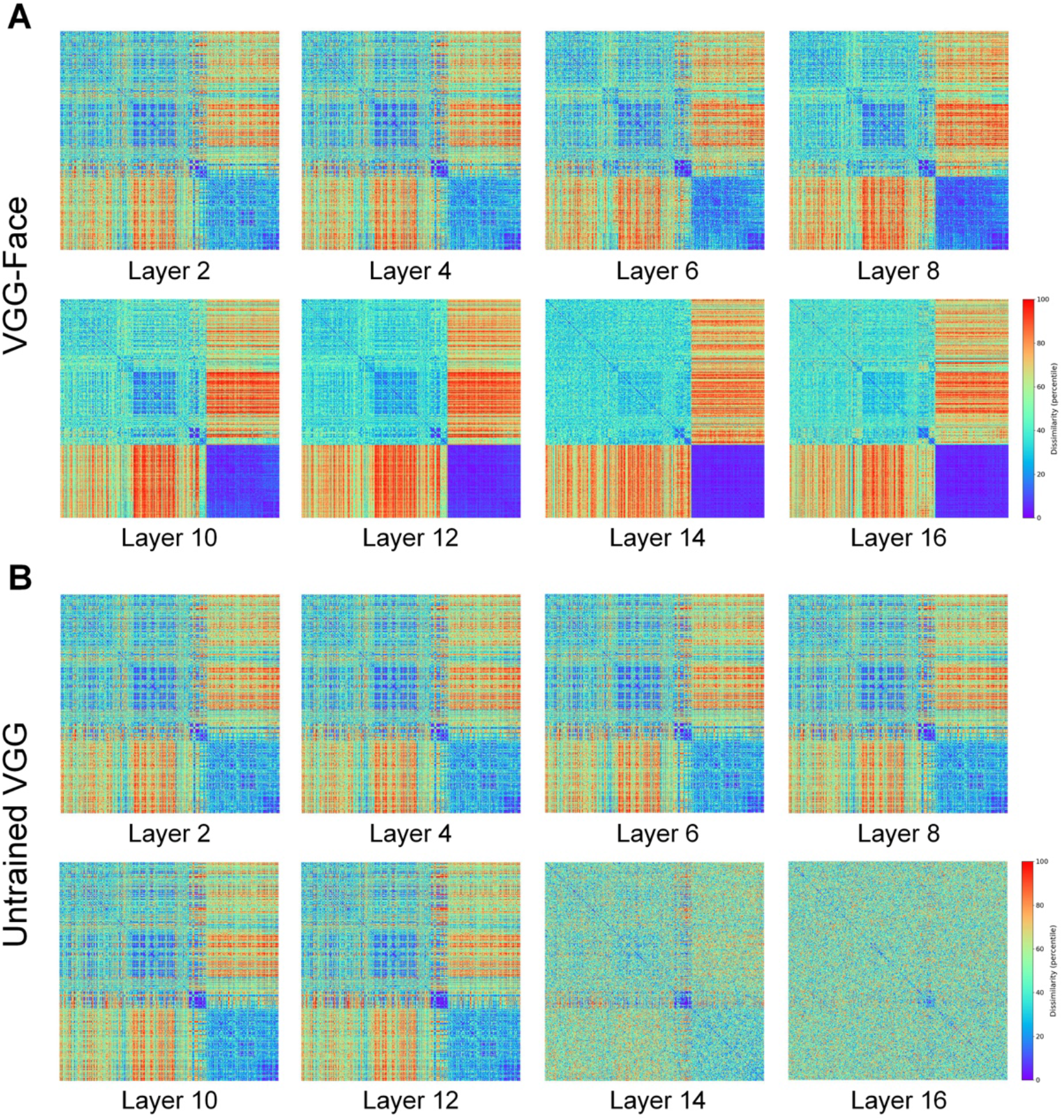
DCNN 450×450 RDMs based on (A) VGG-Face and (B) Untrained VGG.

**Supplementary Figure 3.**
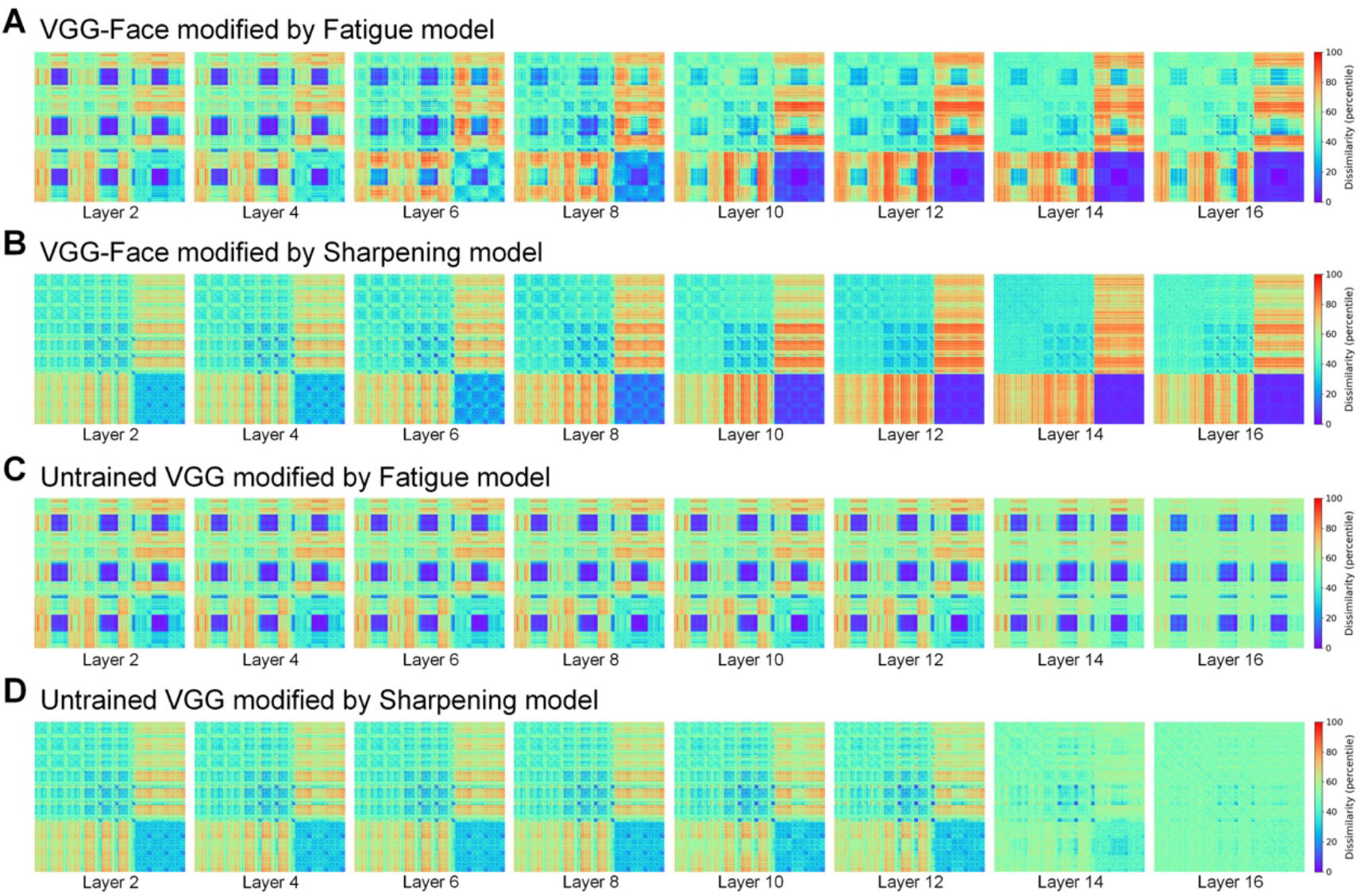
1350×1350 RDMs based on (A) VGG-Face modified by Fatigue model, (B) VGG-Face modified by Sharpening model, (C) untrained VGG modified by Fatigue model, and (D) untrained VGG modified by Sharpening model.

## Notes

### Competing Interest Statement

The authors have declared no competing interest.

### Summary of Updates

manuscript revised

